# Adolescent dopamine neurons represent reward differently during action and state guided learning

**DOI:** 10.1101/2021.07.05.451195

**Authors:** Aqilah M. McCane, Meredyth A. Wegener, Mojdeh Faraji, Maria T. Rivera Garcia, Kathryn Wallin-Miller, Vincent D. Costa, Bita Moghaddam

## Abstract

The neuronal underpinning of learning cause-and-effect associations in the adolescent brain remains poorly understood. Two fundamental forms of associative learning are Pavlovian (classical) conditioning, where a stimulus is followed by an outcome, and operant (instrumental) conditioning, where outcome is contingent on action execution. Both forms of learning, when associated with a rewarding outcome, rely on midbrain dopamine neurons in the ventral tegmental area (VTA) and substantia nigra (SN). We find that in adolescent male rats, reward-guided associative learning is encoded differently by midbrain dopamine neurons in each conditioning paradigm. Whereas simultaneously recorded VTA and SN adult neurons have a similar phasic response to reward delivery during both forms of conditioning, adolescent neurons display a muted reward response during operant but a profoundly larger reward response during Pavlovian conditioning suggesting that adolescent neurons assign a different value to reward when it is not gated by action. The learning rate of adolescents and adults during both forms of conditioning was similar further supporting the notion that differences in reward response in each paradigm are due to differences in motivation and independent of state versus action value learning. Static characteristics of dopamine neurons such as dopamine cell number and size were similar in the VTA and SN but there were age differences in baseline firing rate, stimulated release and correlated spike activity suggesting that differences in reward responsiveness by adolescent dopamine neurons are not due to differences in intrinsic properties of these neurons but engagement of different networks.

## INTRODUCTION

Understanding how motivated behavioral states are encoded by the adolescent brain is critical for detection and prevention of brain disorders and reckless behaviors that emerge in this developmental stage. These include, but are not limited to suicide attempts, addiction, mood disorders, and schizophrenia. What about the adolescent neural processing of motivated behavior predisposes them to these conditions? This is a question we are poorly equipped to answer because much of data on neuronal representation of mental processes related to the operation of the motivational systems were generated using adult animal models (Robbins and Everitt, 1996; Dayan and Balleine, 2002; Berridge, 2004; Schultz, 2010; Flagel et al., 2011). These include data related to incentive motivation, reinforcement learning and decision making that factor prominently in potential models and influential theories that attempt to explain adolescent vulnerabilities and reckless behaviors (Ernst et al., 2011; Luciana and Collins, 2012; Naneix et al., 2012; Casey, 2015; Larsen and Luna, 2018; Hauser et al., 2019).

Motivated behavior, rudimentarily defined as an action taken toward an expected outcome, is constrained by learning. The organism can only be motivated about an outcome if it has learned that the outcome may be a consequence of an action or a context. Thus, the neuronal basis of adolescent motivated behavior is guided by the previously learned cause-and-effect associations. Two fundamental and complementary forms of associative learning are Pavlovian (classical) and operant (instrumental) conditioning (Dickinson, 1981; Fanselow and Wassum, 2015; Corbit and Balleine, 2016). Pavlovian conditioning involves learning that a particular stimulus (conditioned stimulus, CS) in the environment predicts the occurrence of an outcome (unconditioned stimulus, US), independent of any action taken. Operant conditioning involves learning that a particular action by the organism leads to the occurrence of a reinforcing outcome. Each learning process can be described by temporal difference learning algorithms that differentiate between state and action values (Averbeck and Costa, 2017). State values are defined by the information that predicts upcoming rewards whereas actions can take on different values depending on the state in which they are enacted.

Neuronal networks and circuits that contribute to these forms of learning are multi-dimensional, and involve multiple and distinct brain regions (Maren, 2001; Fanselow and Wassum, 2015; Corbit and Balleine, 2016; O’Doherty, 2016). Both forms of conditioning, however, involve midbrain dopamine neurons. In the adult brain, dopamine neurons in the ventral tegmental area (VTA), which project primarily to ventral (limbic) striatal regions and represent state and action values as well as reward prediction errors, are important for reward signaling during both types of conditioning (Schultz, 1998; Pessiglione et al., 2006; Keiflin et al., 2019). Emerging data suggest that adult dopamine neurons in another midbrain region, the substantia nigra (SN) are also involved in processing reward-related learning (Coddington and Dudman, 2018; Keiflin et al., 2019; van Zessen et al., 2021). Dopamine neurons in SN primarily project to the dorsal striatum. Notably, adolescent rodents exhibit large reward-related firing in the dorsal striatum compared to adults (Sturman and Moghaddam, 2012b), suggesting a nigrostriatal bias in encoding reinforcement learning.

To better understand if and how adolescent VTA and SN neurons encode cause-and-effect relationships differently than adults, we recorded from these regions simultaneously during Pavlovian and operant conditioning in both age groups. The US in the Pavlovian task and the action-led outcome in operant task involved the delivery of an identical food reward allowing for comparison of the operational aspects of these learning paradigms independent of the expected outcome. The data across these two learning tasks suggest that despite the same learning rate as adults, adolescents employ different patterns of dopamine neuron activation to reach the same reward endpoint.

## RESULTS

### Learning rates and behavioral performance are similar in both adults and adolescents

Adult and adolescent rats were trained in Pavlovian or operant conditioning as we recorded from VTA and SN neurons. The learning paradigms were designed so that, while operationally distinct, used the same behavioral apparatus and resulted in the delivery of the same reward (a sugar pellet) in each trial (Figure 1A). We assessed learning during consecutive operant (n=16 adults, n=6 adolescent) and Pavlovian (n=11 adults, n=4 adolescents) conditioning sessions by computing the latency to retrieve the sugar pellet that served as the US after CS termination in Pavlovian sessions, the latency to nose-poke into the lit port after CS onset in operant sessions (Figure 1B) and the total number of trials completed (Figure 1C). Age does not influence latency to retrieve in Pavlovian conditioned animals (two-way repeated measures (RM) ANOVA, main effect of age, F(1,15)=0.14,p=0.72) and both age groups decrease their latencies over session (two-way RMANOVA, main effect of session: F(5,62)=6.92,p<0.0001). In operant conditioning, both adolescents and adults show a decrease in their latency to poke over sessions (two-way RMANOVA, main effect of session: F(5,98)=14.80,p<0.0001), but overall adolescents show longer latencies to make a response (two-way RMANOVA, main effect of age: F(1,22)=6.91,p=0.02). Both age groups completed a comparable number of trials during Pavlovian (main effect of age: F(1,34)=0.39,p=0.54) and operant conditioning (main effect of age: F(1,61)=0.02,p=0.88). We then modeled learning based on performance latencies using the Rescorla-Wagner algorithm (Danks, 2003) to estimate individual learning rates across behavioral sessions in both tasks (Figure 1D). Specifically, we estimated the rate at which adolescent and adult rats learned the Pavlovian and operant associations with the predictive cue or action. Learning rates do not support significance of age: F(1,15)=0.003,p=0.96, task: F(1,17)=1.29,p=0.27, or an interaction between age and task: F(1,17)=1.40,p=0.25.

**Figure 1.**
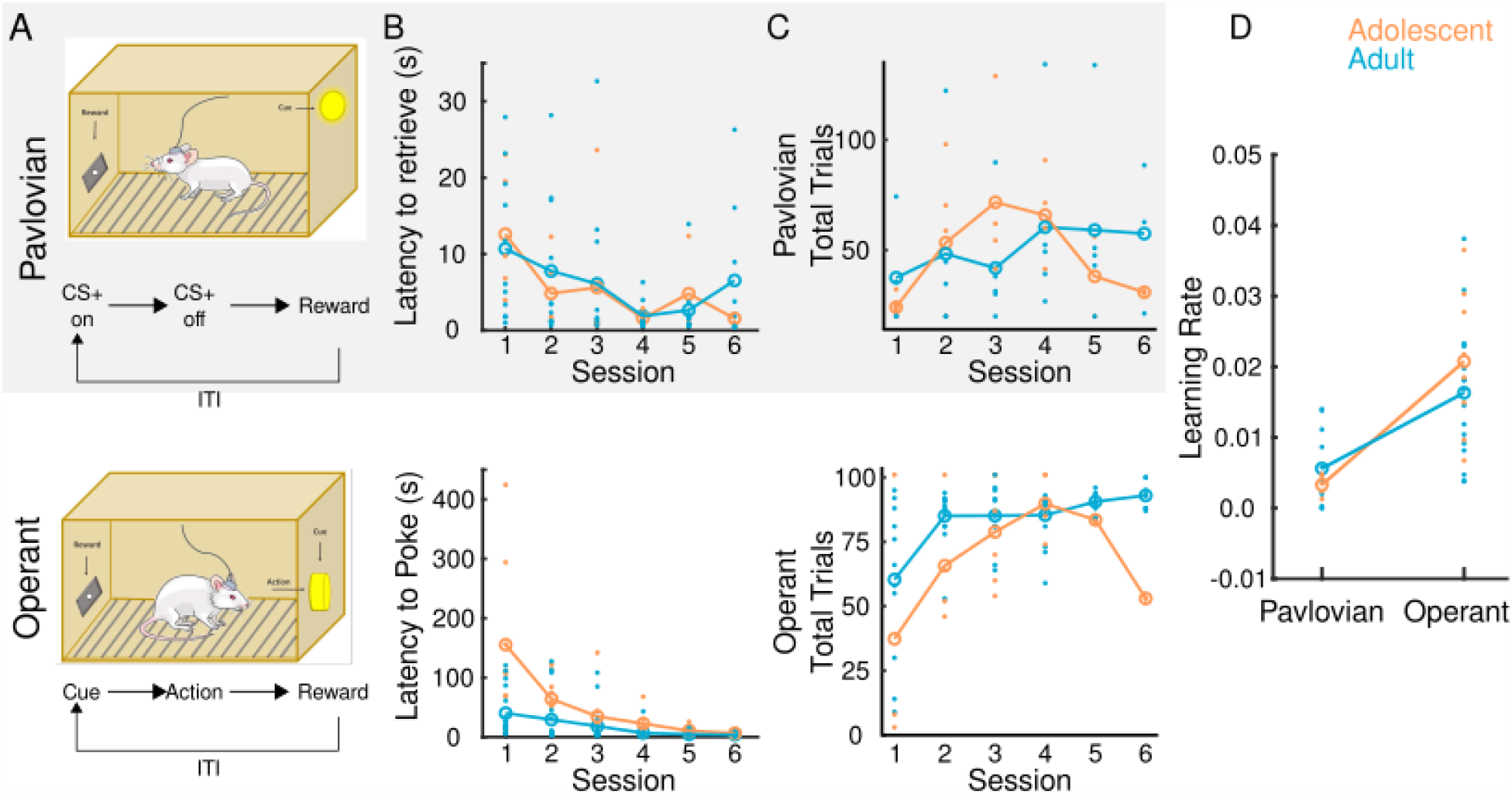
Experimental methods and behavior. (A) Schematic of operant box and behavioral paradigm for Pavlovian (grey) and operant (white) conditioned adolescent (orange) and adult (blue) animals. During Pavlovian conditioning, a CS (a light on the opposite side of the food magazine) was presented for 10 s after which the US (sugar pellet) was delivered. During operant conditioning, a cue (also a light) was presented in a hole located in the opposite side of the food magazine. Executing an action (nosepoke) led to delivery of reward. In both paradigms, once the reward was retrieved, another trial was started after an inter-trial interval. (B) Pavlovian learning was assessed by measuring latency to retrieve a reward and operant learning was assessed by measuring latency to poke across sessions. (C)Total number of trials completed in both tasks was also assessed. (D) Individual learning rate was estimated using Rescorla-Wagner model (eq. 1). Learning rate is scaled from one to zero where one represents instant learning and zero corresponds with no learning. There was no influence of age on learning rate of either task. Similarly, both adolescents and adults complete similar numbers of trials and decrease latency to perform either an action or conditioned approach, collectively demonstrating that both adolescents and adults learn each respective task. Individual points represent data from one animal. Values are means.

### Static and dynamic characteristics of VTA and SN dopamine neurons are similar between adolescents and adults

VTA and SN neurons were classified as putative dopamine based on wave form width greater than 1.2 ms and mean baseline firing rate slower than 12 Hz, as reported previously (Grace and Bunney, 1984; Schultz and Romo, 1987; Kim et al., 2016). This classical approach has been substituted in some recent papers by optogenetic classification of dopamine neurons (Coddington and Dudman, 2018; Mohebi et al., 2019). We and others have observed that the waveform and firing rate of optogenetically identified dopamine neurons is consistent with classically defined criteria (Stauffer et al., 2016; Hughes et al., 2020). Here, we were not able to repeat the opto-tagging characterization because the short time-frame of the adolescent experiments (less than 3 weeks between weaning and the start of recording) does not allow for sufficient viral expression required for optogenetic tagging of dopamine neurons. Instead, we supplemented our waveform and firing rate characterization approach with examining the effect of the dopamine agonist apomorphine on the inter-spike interval of VTA and SN neurons in a separate group of rats (Figure 2A,B). Consistent with previous reports (Guyenet and Aghajanian, 1978; Schultz and Romo, 1987), this treatment increased the inter-spike interval of neurons with dopamine-like waveforms without affecting neurons classified as fast spiking putative non-dopamine neurons (Figure 2 A,B). Only units characterized as putative dopamine neurons (VTA n=272 adults n=241 adolescents, SN n=226 adult n=241 adolescents) were used for further analysis. These neurons displayed canonical phasic response to reward during both Pavlovian (Figure 2C) and operant conditioning (Figure 2D). We next evaluated whether age mediated differences in dopamine neuron characteristics. A two-sample t-test revealed adolescents exhibited faster basal firing rates relative to adults in the SN (t(248.39)=-2.51, p=0.01), but not VTA (t(431.69)=-1.39, p=0.16; Figure 2E). There was no effect of age (n=11 adults, n=14 adolescents) on number of dopamine cells in the SN (t(9)=-0.41,p=0.69) or VTA (t(6)=0.96,p=0.37; Figure 2F). Also, there was no effect of age on dopamine cell size in the SN (t(10)=0.68,p=0.51) or VTA (t(10)=0.14,p=0.89; Figure 2F).

**Figure 2.**
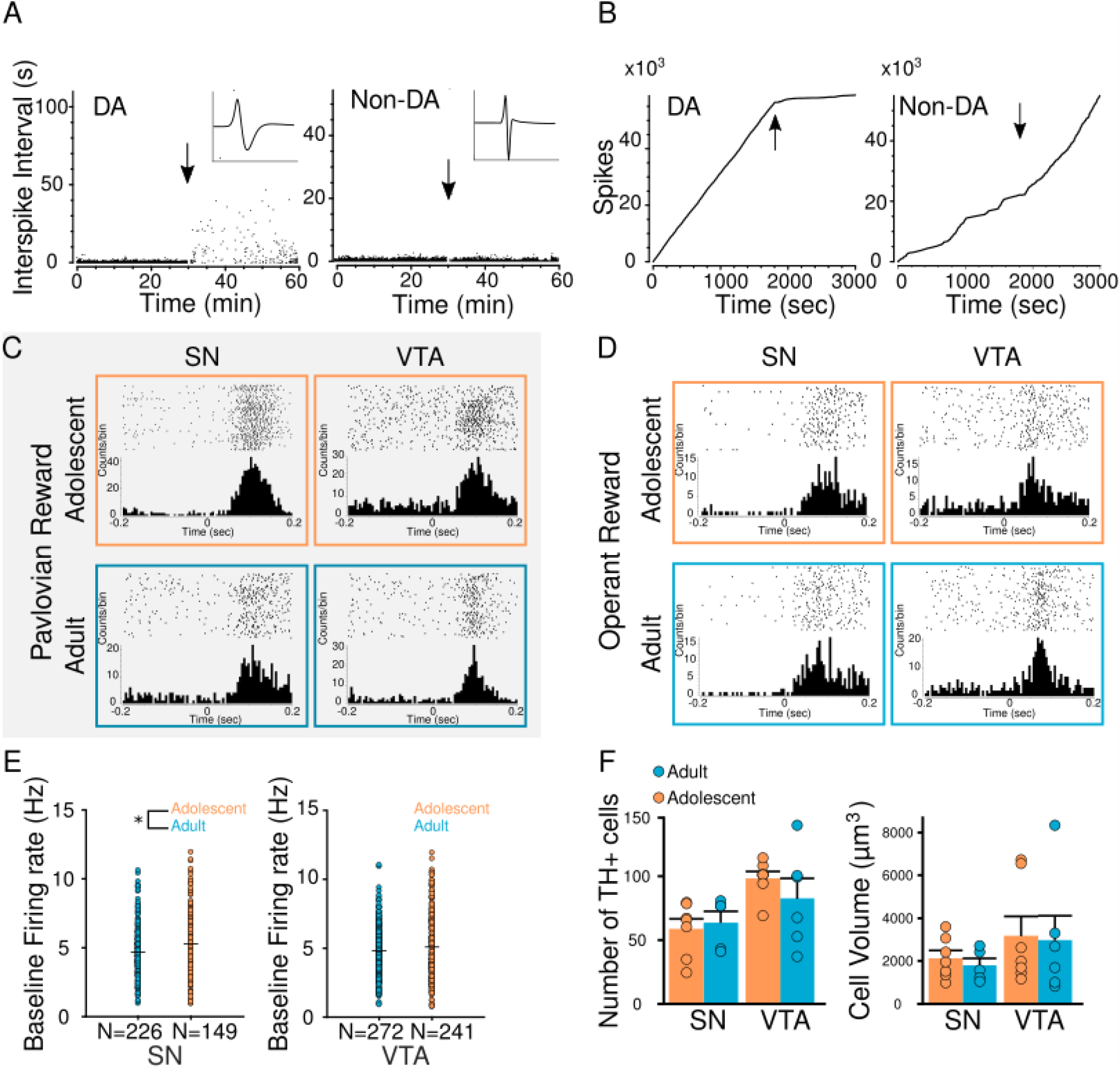
Dopamine neuron classification and characterization. We classified neurons as putative dopamine or non-dopamine based on waveform width and mean baseline firing rate. As a secondary classification metric, we measured the impact of the dopamine agonist apomorphine (0.75 mg/kg) on the inter-spike interval. Arrows indicate drug administration time point. (A, B).Apomorphine increase the inter-spike interval in putative dopamine neurons but has no effect on non-dopamine neurons. Raster plots of individual neuron firing in response to reward (C, D) As expected, neurons classified as dopamine exhibited a phasic response to reward during both Pavlovian (C) and operant (D) conditioning. (E) Baseline firing rate in the VTA in adults and adolescents was similar but adolescents exhibit slightly higher firing rate in the SN. (F) Dopamine cell volume and numbers were similar in the VTA and SN in adolescent (PND 35) and adult rats (PND 75). *p<0.05, t-test. Values are mean ± SEM.

### Session-by-session response of dopamine neurons revealed age- and task specific differences during learning

Figure 3 shows the phasic response to presentation of the CS and US during Pavlovian conditioning. Phasic response to reward in the SN was larger in adolescents compared to adults (main effect of age: F(1,158)=9.39,p=0.003; Figure 3A). A non-significant trend in peak firing rate changes across session was also observed (main effect of session: F(5,158)=2.23,p=0.05). In the VTA, peak firing rate was influenced by both age and session: F(5,227)=2.37,p=0.04; Figure 3B). Age differences in phasic response to reward were most pronounced on session 3 and later. The larger dopamine response in adolescents during and after session 3 was specific to reward and did not generalize to phasic response to the other events including CS initiation. Could differential valuation of the reward *per se* evoke a larger response by dopamine neurons? Session-by-session analysis of the response of dopamine during operant conditioning suggested that this is not case. As this form of conditioning progressed, dopamine neurons response to reward was smaller in adolescents compared to adults in both absolute and normalized levels in SN and VTA (Figure 4). Compared to adolescents, a larger phasic response was observed in adults during reward in the SN (main effect of age: F(1,249)=6.34,p=0.01; Figure 4A) and VTA (main effect of age: F(1,321)=8.32,p=0.004; Figure 4B). There was no effect of session in either brain region (p values >0.05).

**Figure 3.**
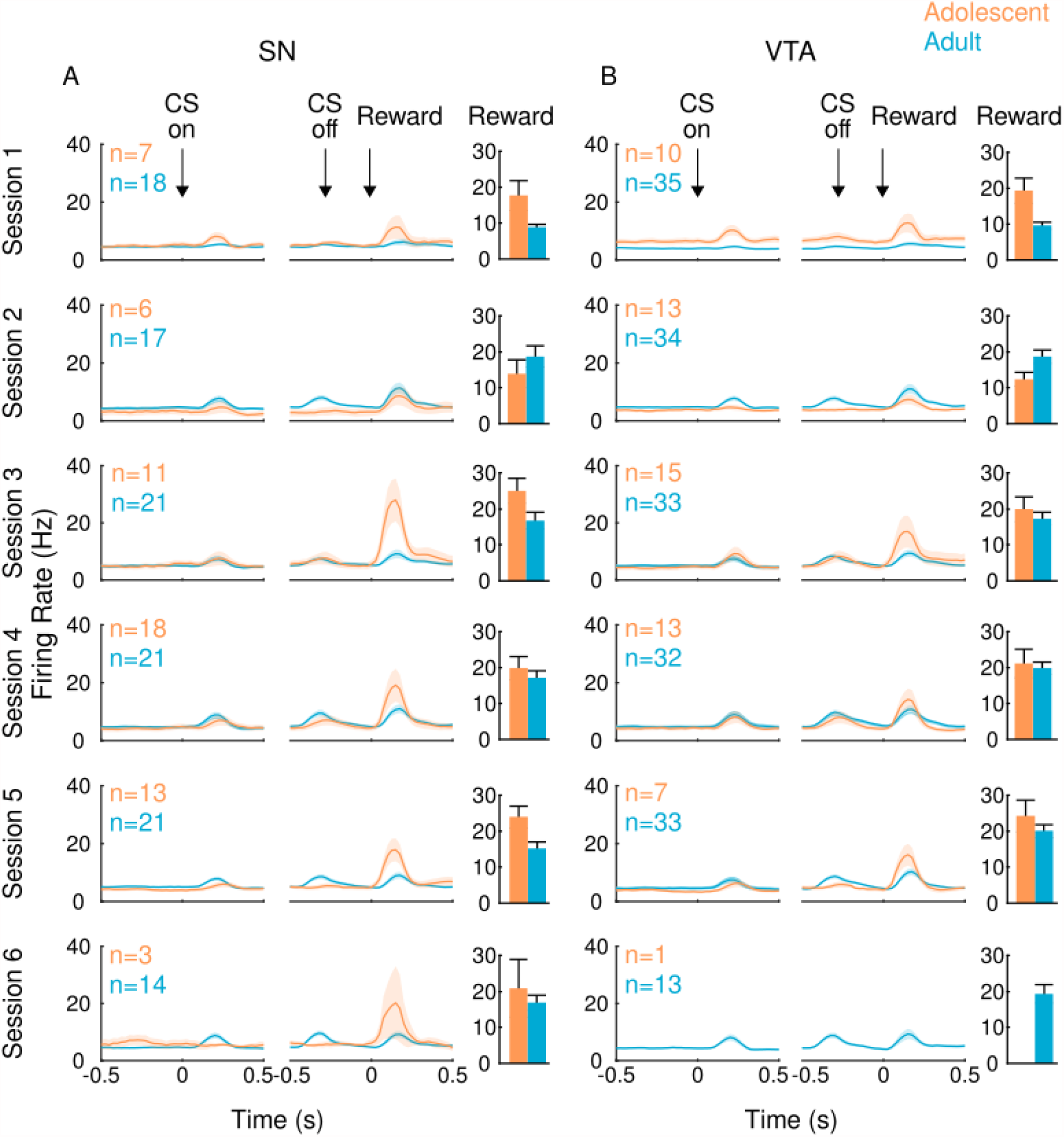
Pavlovian firing rates across session. Session-by session firing rates are in adolescent (orange) and adult (blue) rats in the SN (A) and VTA (B). Bar graphs reflect mean peak phasic response 500 ms after reward. Dopamine neurons recorded in adolescents exhibited a larger response to reward receipt than dopamine neuron responses recorded in adults. This effect was more pronounced among dopamine neurons recorded in the SN versus the VTA. Values are mean ± SEM.

**Figure 4.**
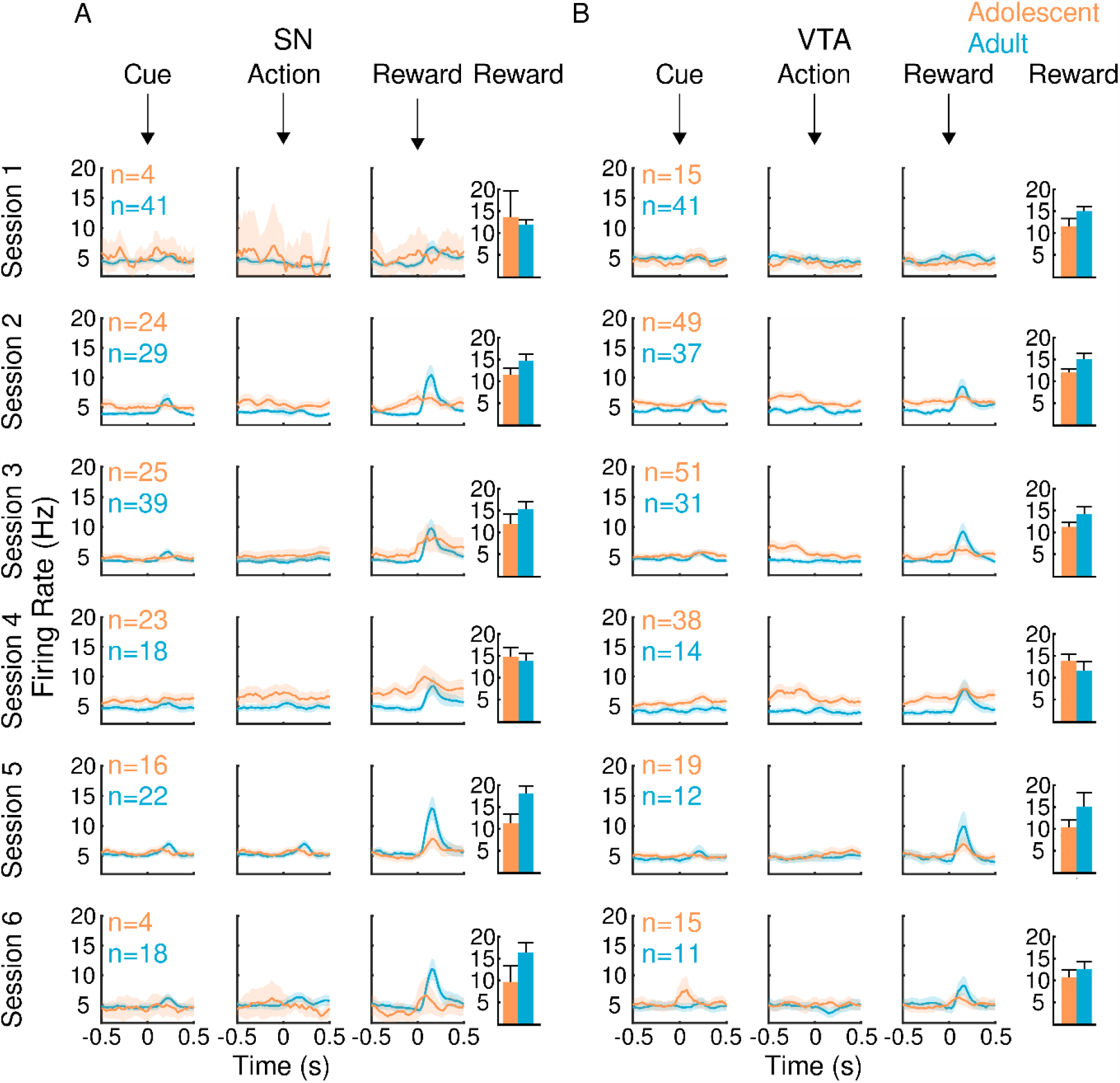
Operant firing rates across session. Session-by session firing rates are in adolescent (orange) and adult (blue) rats in the SN (A) and VTA (B). Bar graphs reflect mean peak phasic response 500 ms after reward. Dopamine neurons recorded in adults exhibited a larger response to reward receipt than dopamine neuron responses recorded in adolescents. Values are mean ± SEM.

Comparison of adult and adolescent phasic response to reward and other key events during Pavlovian and operant conditioning was made by considering the normalized response across sessions (3-6) in which performance in both tasks was stable. We first determined whether significant events such as CS or reward presentation evoked a significant change in firing rate. Firing rate was significantly altered by CS presentation in all animals (main effect of epoch: adolescent SN: F(2,82)=7.62,p=0.0009; adolescent VTA: F(2,68)=7.93,p=0.0008; adult SN: F(2,168)=65.19,p<0.01; adult VTA: F(2,194)=68.86,p<0.001) where a robust phasic response to presentation of the CS was observed in both the VTA and SN of both age groups (Figure 5A). During cue presentation in operant conditioning, a main effect of epoch was observed in the SN of adults (F(2,186)=15.15,p<0.001) but not adolescents (F(2,114)=2.076,p=0.13) and the VTA of both age groups (adult: F(2,68)=11.95,p<0.001; adolescents: F(2,166)=4.84,p=0.009;Figure 5B). A phasic response to Pavlovian reward delivery was observed in both adolescents and adults in both the SN (adolescents: F(2,82)=21.56,p<0.001;adults: F(2,168)=14.29,p<0.001) and VTA (adolescents: F(2,68)=9.64,p=0.0002 adults: F(2,194)=39.78,p<0.001; Figure 5C). In the operant group, a main effect of epoch was observed in both brain regions in adults (SN: F(2,186)=19.26,p<0.001; VTA :F(2,126)=12.63,p<0.001) and the SN of adolescents (F(2,114)=5.98,p=0.003; Figure 5D).

**Figure 5.**
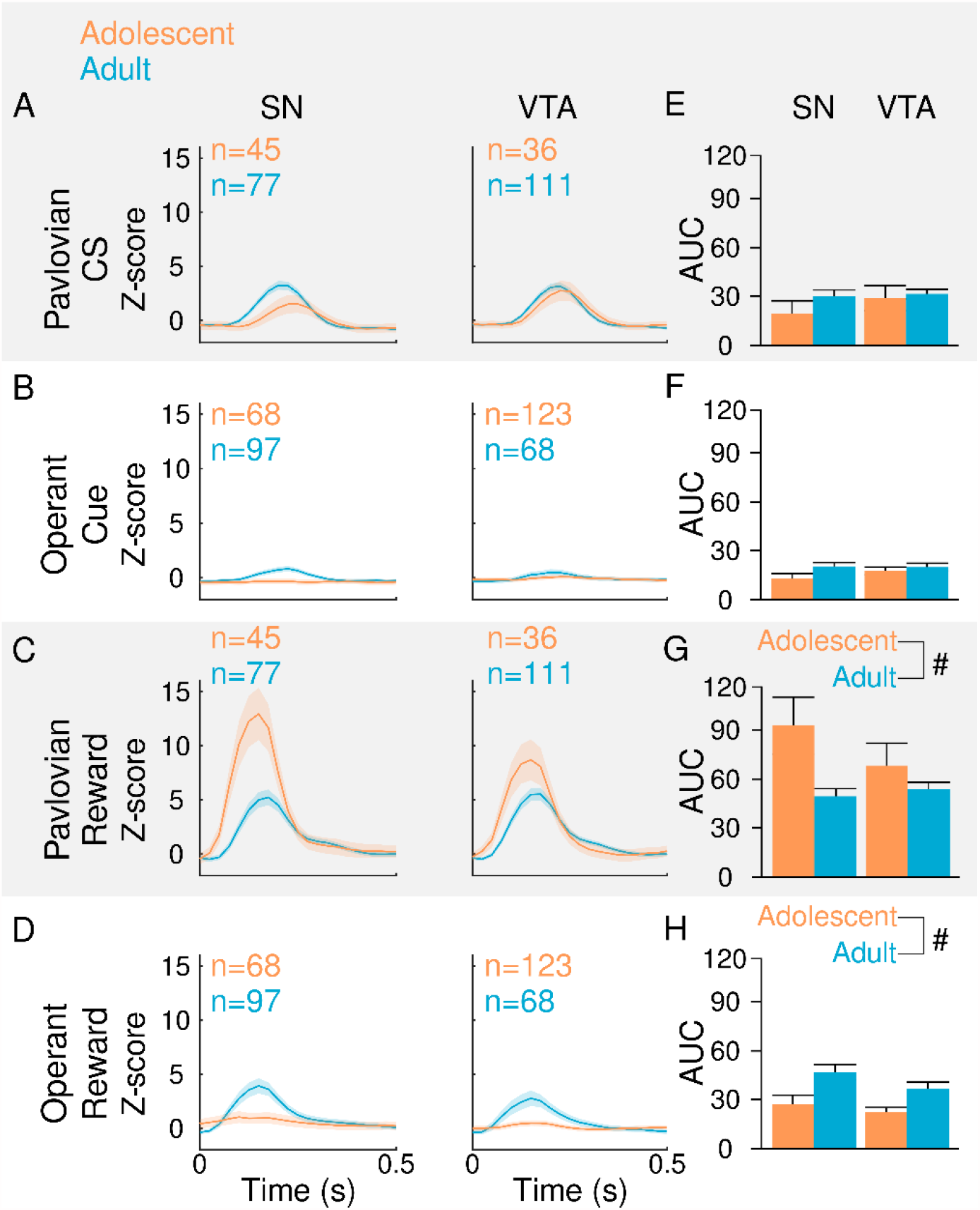
Averaged firing rates around salient events. Adolescent (orange) and adult (blue) data was combined across sessions 3-6, after both tasks were learned. To determine whether the phasic response following behavioral events of interest was significantly different from basal activity, data were divided into 500ms epochs before, during and after each event, and RMANOVAs were performed in data stratified by brain region and age. Phasic responses to CS presentation in Pavlovian conditioning (A) and operant conditioning (B) were comparable in both ages. Compared to adults, adolescents exhibited a larger phasic response to reward during Pavlovian conditioning in both the SN and VTA (C, G). In contrast, adults exhibited a pronounced phasic response to reward during operant conditioning in both the SN and VTA while the phasic response in adolescents was attenuated, relative to adults and only observed in the SN (D, H). #p<0.05, main effect of age. Values are mean ± SEM.

We next compared firing rate between behavioral groups, brain regions and age groups. Area under the curve (AUC) was computed for the 500ms epoch following each event. Group differences were then assessed by ANOVA with Bonferroni corrected post hoc tests performed as necessary. During Pavlovian CS presentation, there was no effect of age (main effect of age: F(1,710)=1.16,p=0.28) or brain region on AUC firing rate (main effect of brain region: F(1,710)=0.30,p=0.59; Figure 5E). During cue presentation in operant conditioned rats, there was no effect of age (main effect of age: F(1,491)=2.93,p=0.09) or brain region on firing rate AUC (main effect of brain region: F(1,491)=0.71,p=0.40; Figure 5F). During reward delivery, overall firing rate was greater in the Pavlovian group, compared to the operant group (main effect of task: F(1,1146)=66.42,p=1.03×10^−15^). Data were therefore next stratified by behavioral task. In the Pavlovian group adolescents exhibited greater firing rate during reward (main effect of age: F(1,660)=10.76,p=0.001; Figure 5G). There was no difference between brain regions in the Pavlovian animals (main effect of brain: F(1,660)=0.0,p=0.98). In contrast, adults in the operant group exhibited greater firing rate during reward delivery (main effect of age: F(1,486)=18.95,p=1.64×10^−5^), with a non-significant trend between brain region differences observed (main effect of brain: F(1,486)=3.09,p=0.07; Figure 5H). In summary, there was no effect of age on CS presentation in either paradigm, but adolescents exhibited a larger phasic response to reward during Pavlovian conditioning in both the SN and VTA compared to adults, while adults exhibited a more pronounced phasic response to reward during operant conditioning, compared to adolescents. This analysis further established that the same reward achieved as a US, as opposed to obtained after an action, selectively produces a larger response in adolescents.

### Adolescents VTA and SN neurons exhibit different correlated activity to reward in different conditioning paradigms

Neuronal representation of behavioral events can be distributed across populations of neurons (Cohen et al., 2012).To assess population dynamics in response to reward during learning in SN and VTA of adult and adolescents, we computed spike correlation in simultaneously recorded neurons (Cohen and Kohn, 2011; Kim et al., 2012). Data were stratified by brain region and behavioral task, and two-way repeated measures ANOVAs were performed to determine whether spike correlation ratios after stimulus (cue or CS) presentation changed across session and whether this effect was influenced by age group. In all groups, the main effect of sessions was not significant (p values>0.05). Adolescents in the Pavlovian group, exhibited more correlated activity than adults in the SN (main effect of age F(1,22)=42.34,p=1.52×10^−6^) and the VTA (main effect of age (F,23)=23.39,p=7.01×10^−5^; Figure 6A). Similarly, in the operant group, a main effect of age was observed in the SN (F(1,46)=20.96,p=3.56×10^−5^) and VTA (F(1,68)=56.08,p=1.86×10^−10^; Figure 6B). Differences in population response to conditioned stimuli may reflect differences in encoding efficiency or the amount of information encoded by that population and received by downstream networks.

**Figure 6.**
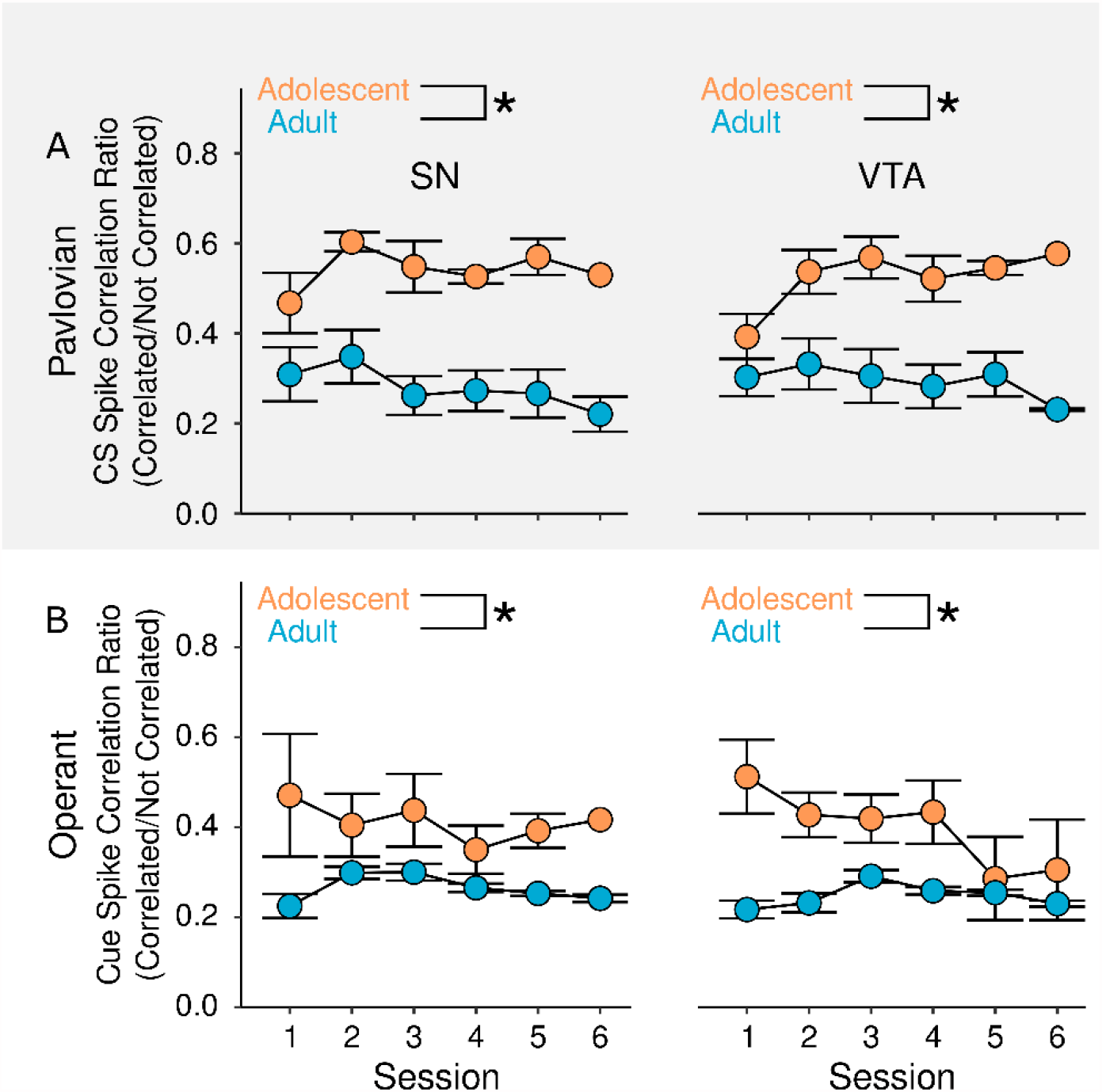
Spike correlation ratios (correlated / non-correlated neuron pairs) during Pavlovian and operant conditioning. The higher the ratio, the greater the number of correlated pairs, or the smaller the variance in population dynamics. Conversely, a lower ratio indicates fewer correlated pairs and a more heterogeneous population dynamic. In comparing coordinated activity in adults and adolescents during stimulus presentation, we observed an age-mediated different pattern of correlated activity in both Pavlovian (A) and operant (B) conditioning. *p<0.05, main effect of age. Values are mean ± SEM.

### Terminal dopamine release in response of activation of dopamine neurons is muted in adolescents

Motivated actions are mediated by dopamine release from the terminals. Mechanisms that govern dopamine release and volume transmission may be different in adults and adolescents (Robinson et al., 2011; Pitts et al., 2020). Our observation of increased dopamine neuron phasic response to reward in Pavlovian conditioning but decreased, phasic response in operant conditioning could be functionally amplified or muted if dopamine release is different in response to similar phasic activation of these neurons. We, therefore, determined if the same pattern of activation of dopamine neurons in adults (N=34) and adolescents (N=31) produces similar increase in terminal release. Dopamine efflux was measured in nucleus accumbens and dorsal striatum in response to different patterns of stimulation which mimicked different patterns of dopamine neuron activation (Lohani et al., 2019). These two regions have been implicated in both forms of conditioning (O’Doherty et al., 2004; Day and Carelli, 2007; Corbit and Janak, 2010). Phasic burst stimulation (20 pulses at 100 Hz) altered dopamine release in the DS (main effect of time: F(9,135)=21.14,p<2×10^−16^) but was not influenced by age group (main effect of age: F(1,15)=0.26,p=0.62; Figure 7B). Phasic burst stimulation also elicited an increase of dopamine in the NA which was influenced by rodent age (age × time interaction: F(9,135)=2.81,p=0.005). Specifically, both adults and adolescents exhibited an increase in dopamine at samples 5,6,7 (Dunnett’s post hoc, p values<0.05). However, this increase was greater in adults (main effect of age: F(1,15)=4.78,p=0.04). Phasic sustained (100 pulses at 20 Hz) stimulation of the VTA produced a mild increase in dopamine levels in the DS (two-way RMANOVA, main effect of time: F(9,108)=18.16, p<2×10^−16^) and the NA (F(9,135)=17.77,p<2×10^−16^; Figure 7C), which was similar between adolescents and adults (main effect of age, p values >0.05). With regard to locomotor activity, phasic burst stimulation increased the number of fine movements (main effect of time:F(10,160)=6.34,p=3.59×10^−10^) but was not influenced by age group (main effect of age:F(1,16)=3.03,p=0.10;Figure 7D). Sustained stimulation did not increase locomotion in any of the groups (two-way RMANOVA, main effect of time: F(10,132)=1.17,p=0.31; Figure 7E). In summary, the general trend we observed was reduced dopamine release in adolescents with the most robust effect observed in the nucleus accumbens after burst activation of dopamine neurons.

**Figure 7.**
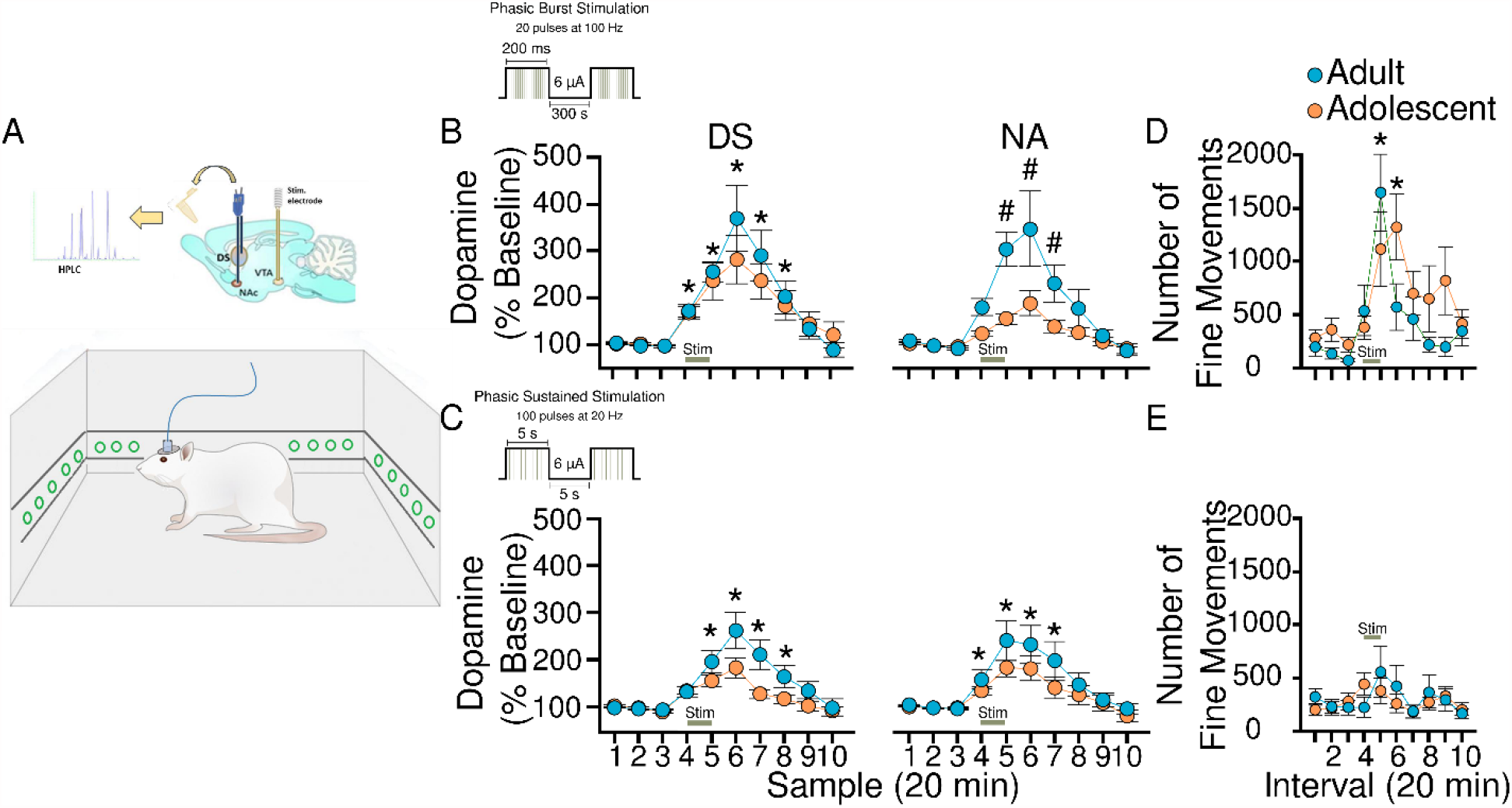
Microdialysis measurement of evoked dopamine release. (A) Schematic illustration of VTA electrical stimulation and simultaneous measurement of dopamine dialysates in the dorsal striatum (DS) or nucleus accumbens (NA) and locomotor activity in freely moving in adolescent (orange) or adult (blue) rats. (B) Phasic burst stimulation was associated with increased dopamine release in the DS and NA in both adults and adolescents but in the NA the effect of stimulation was more pronounced in adults. (C) Phasic sustained stimulation resulted in an increase in dopamine release in both the DS and NA, comparable in both age groups. Phasic burst stimulation increased the number of fine motor movements in both age groups (D), while sustained stimulation had no effect on locomotor activity (E). Data are expressed as mean ± SEM. *p<0.05 days different from baseline, Dunnett’s post hoc. # p<0.05, main effect of age.

## DISCUSSION

Reckless behavior and impulsive decision making by adolescents suggest that motivated behavioral states are encoded differently by the adolescent brain (Simon and Moghaddam, 2015). Motivated behavior follows learning of cause-and-effect associations in the environment. Here we sought to understand if the learning of these associations, and response of dopamine neurons in VTA and SN during learning, differ in adolescents as compared to adults. We focused this work on comparing two elementary forms of associative learning: Pavlovian and operant conditioning. In both conditioning paradigms, we used the same reward as the outcome. This ensured that the rewarding value of the outcome was identical and, therefore, behavioral or neural differences in the two paradigms were due to operational differences in the means to reach the outcome. We find that while learning rate is similar in both ages, adolescent dopamine neurons encode reward differently depending on the cause-and-effect relationship of the means to receive that reward. Compared to adults, reward contingent on action led to a muted response whereas reward that was not gated by action produced an augmented response, suggesting adolescent dopamine neurons assign a high value to rewards that are made available independent of actions.

### Learning rate and response rate in Pavlovian and operant conditioning in adolescents and adults

Pavlovian conditioning involves learning that occurrence of a stimulus in the environment predicts the occurrence of an outcome, independent of taking any particular action. In the parlance of reinforcement learning, Pavlovian learning amounts to learning the value of state that occurs when the conditioned stimulus is presented. We found that the learning rate of adolescents is not different from adults during Pavlovian conditioning suggesting that state value representations are intact in adolescents.

During operant conditioning, the rats can control how and when a reinforcing event occurs by deciding to execute an action in a particular state. The action value of making a nosepoke comes to define the ultimate value of the state in which that action is executed. We found that while adolescents exhibited longer latencies to execute an action, they displayed similar learning rate suggesting that they can learn contextualized action values similar to adults. The longer latencies to make an operant response, despite having learned the action-reward association, suggest lower motivation and slower capacity to update action values after action execution.

### Adolescent VTA and SN neurons are engaged differently to reach the same behavioral endpoints as adults

Dopamine neurons in the VTA and SN have been implicated in operant and Pavlovian conditioning (Schultz, 1998; Dalley et al., 2002; Parkinson et al., 2002; Haruno and Kawato, 2006; Lex and Hauber, 2010; Coddington and Dudman, 2018; Keiflin et al., 2019; van Zessen et al., 2021). While much of the learning literature has focused on dopamine neurons in the VTA, multiple studies indicated that SN neurons also generate reward related signals (Coddington and Dudman, 2018; Saunders et al., 2018). In adult rats, we find that neurons in VTA and SN have a near identical magnitude of response to reward during either conditioning paradigm. Both neuron groups displayed similar phasic responses to operant cue and Pavlovian CS. Moreover, phasic response of adult VTA and SN neurons to reward delivery was similar regardless of whether it was delivered as US or in response to action execution.

Adolescent neurons, however, had a different response to reward depending on the conditioning paradigm and contingencies that led to reward delivery. In operant conditioning, both VTA and SN cells displayed a smaller phasic response to reward compared to adults. The phasic response of adolescents to the operant cue was equally muted consistent with previous findings (Kim et al., 2016).The lower dopamine activation in adolescents may provide a mechanism for our observation that the latency of action to reward retrieval was longer in adolescents during operant conditioning and is consistent with dopamine’s role in motivation for effort-based behavior (Salamone et al., 2018).

In contrast to the muted dopamine reward response in operant conditioning, adolescents had a robust reward response during Pavlovian conditioning in both regions with the SN response being slightly larger than VTA. Thus, adolescent dopamine neurons may assign higher value to a given reward when it is obtained independent of action.

What could be the potential mechanism for the difference in contingency-dependent signaling of dopamine neurons in response to the same reward? Importantly, there was no age difference in dopamine cell number or size. Thus, age differences in reward-evoked activity cannot be explained by static dopamine neuron characteristics. This difference is also unlikely to be due to phasic vs tonic firing activity relationship (i.e. lower tonic activity promoting higher or lower phasic response as predicted by Lucia and Collins, 2012). While, dopamine neurons in SN had a higher tonic firing rate, these neurons had a muted phasic reward response during operant conditioning and elevated phasic response during Pavlovian conditioning. Therefore, the age-specific differences in VTA and SN activity reported here are likely due to different networks driving dopamine neurons during reward. Consistent with this notion, we observed age-specific changes in spike correlation. This measure reflects the strength of information provided by a population of neurons to their target regions (Cohen and Kohn, 2011) and thus may be an index of functional connectivity among networks of neurons. While the difference we observe was not task-specific, it suggest that activation of distinct networks are involved in phasic activation of VTA and SN neurons in adults and adolescents. A relatively large literature has, in fact, implicated distinct striatal and cortical circuity in operant and Pavlovian conditioning (Cardinal et al., 2002; Shiflett and Balleine, 2011; Peak et al., 2019). In particular, operant conditioning relies on participation of prefrontal cortical regions including the orbitofrontal cortex (OFC) and dorsal striatal regions (McDannald et al., 2005; Yin et al., 2005). These cortical and striatal regions, which have reciprocal connections with dopamine cells, are undergoing maturation during adolescence (Huttenlocher, 1979; Lebel et al., 2008). The OFC and dorsal striatal neurons of adolescents display a large excitatory phasic response to reward during operant conditioning (Sturman and Moghaddam, 2011, 2012b) compared to adults. While future work is needed to delineate age-related differences in circuits that serve associative learning, it is tempting to speculate that weaker dopamine reward response during operant conditioning reduces the postsynaptic dopamine-mediated inhibition on target regions, causing an exaggerated excitatory response in dorsal striatum and OFC. Thus, a muted dopamine neuron response to reward during operant conditioning in adolescents, may lead to increased engagement of DS and OFC, two regions that have been strongly implicated in habit learning (Gremel and Costa, 2013; Barker et al., 2015).

### Adolescents exhibit nigrostriatal bias

An age-dependent difference in overall baseline firing rate was detected in the SN and not the VTA suggesting that, in behaviorally engaged animals, tonic activity of SN neurons is higher in adolescents. Additionally, phasic response to Pavlovian reward in the SN was larger compared to the VTA response and muted response of dopamine release in response to investigator-administered stimulation was only observed in ventral and not dorsal striatum. Dopamine neurons in the SN preferentially innervate the dorsal striatum whereas VTA DA neurons project to the ventral striatum (Haber, 2016). Age-mediated differences in SN and VTA activity together with differences in dorsal and ventral striatal release suggest that adolescents more prominently engage the nigro-dorsal-striatal as opposed to the VTA-accumbal pathway. Interestingly the increase in reward-related firing in the dorsal striatum of adolescents is not observed in ventral striatum (Sturman and Moghaddam, 2012b), further suggesting a bias towards use of nigrostriatal systems in reward processing in adolescents.

### Conclusions and potential interpretations

Adolescent learning rate is similar to adults during Pavlovian and operant conditioning paradigms indicating that their capacity to learn state value representations and contextualized action values are similar to adults. During learning, however, adolescent VTA and SN dopamine neurons exhibited different pattern of paradigm-specific phasic response to reward. Whereas adult neurons responded similarly to reward in both paradigms, adolescent neurons had a larger response to reward delivered as a Pavlovian unconditioned stimulus, and muted response when the same reward was delivered after an action. This observation has two implications. First, it invites the field to rethink influential theories that propose blanket dopamine hyper- or hypo-responsiveness to reward to explain adolescent behavior (Spear, 2000; Ernst and Luciana, 2015; Luna et al., 2015). Our findings clearly demonstrate that while there is an age-related difference in dopamine neuron response to reward, this difference is not uniform and is guided by network processes that differentiate between state and action values. Second, our findings may have evolutionary significance. Pavlovian associations allow organisms to make predictions about occurrence of critical events such as reward availability. Compared to operant conditioning, which may model aspects of foraging behavior, assigning higher motivational value to unconditioned reward availability associated with Pavlovian conditioning can be advantageous because it helps conserve energy. On the other hand, the lower response of dopamine neurons to reward delivery during operant conditioning may be consistent with adolescents being resistant to reward devaluation in instrumental responding (Serlin and Torregrossa, 2015; Marshall et al., 2020; Towner et al., 2020) because it suggests that adolescents assign lower value to the reward, as opposed to cue-action component of this form of conditioning. This may also be evolutionary advantageous because it allows adolescents to explore (Gopnik et al., 2021) and persist in goal-directed actions and exploratory behavior in the absence reward availability.

## METHODS

### Subjects

Experiments were started at the University of Pittsburgh and completed Oregon Health and Science University. Subjects for all experiments were male Sprague-Dawley rats (Harlan, Frederick, MD; Charles River Laboratories) housed in a humidity and temperature-controlled conditions using a 12-hour reverse light/dark cycle with lights off at 8:00 or 9:00 am. All procedures were approved by either the University of Pittsburgh Institutional Animal Care and Use Committee or the Oregon Health and Science University Institute Animal Care and Use Committee and were in accordance with the National Institutes of Health Guidelines for the Care and Use of Laboratory Animals.

### Conditioning Behavior

Operant chamber (Coulbourn Instruments, Allentown, PA) equipped with a food trough and reward magazine opposite a nose-poke port with a cue light and infrared photo-detector unit, and a tone-generating speaker were used. One day before the start of habituation animals were food restricted to 85% of their weight. After 2-3 days of habituation, where animals learned to retrieve reward from the food magazine, they completed either Pavlovian or operant conditioning. In the Pavlovian task, a light cue (CS) was presented for 10 s on the wall opposite of the food trough. 500 milliseconds (ms) after the termination of the CS, a sugar pellet reward (45 mg sugar pellet, Bio-Serv, Frenchtown, NJ) was delivered. Reward retrieval was followed by a variable (9-12s) inter-trial interval. Each conditioning session consisted of 100 trials. In the operant task, rats were trained to execute an action (nose poke into the lit port) to earn a single sugar pellet reward on a fixed ratio one schedule as described previously (Sturman and Moghaddam, 2012a). Immediately after the action execution, the cue light was extinguished and the reward was delivered after a 1 s delay. Reward collection was followed by 10 s inter-trial-interval and initiation of the next trial. For each trial, the cue light remained illuminated until the rat responded. Each session lasted 45 min or 100 trials.

### Learning Rate

Animal learning behavior was simulated using Rescorla-Wagner model (Danks, 2003) as follows: (Eq. 1),

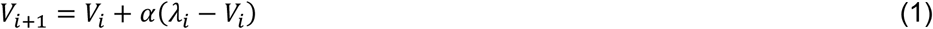

where *V* is the associative value of cue to reward, α represents learning rate, and λ is the maximum reward available. Learning rate is bounded by 0 and 1, where 0 indicates no learning and 1 indicates instant learning, and λ is 0 or 1 based on animal behavior in trial (explained below). Associative value *V* is set to 0 at the beginning of session 1, and as learning takes place this value updates using eq. 1.

Latency of animals to retrieve the reward after delivery (in Pavlovian task), or to poke after cue (in operant task) was used as an indicator of reinforcement learning stage. A latency threshold of 5 seconds was established to distinguish between random and aimed retrievals/pokes, so that the value of a retrieval/poke is one (λ = 1) if it happens within five seconds from delivery/cue, and zero (λ = 0) if after five seconds.

Task sessions were stitched together to allow a better fit for the gradual decrease in latency, and a ten-trial moving average window was applied to smooth the latency data. A mapping function (Eq. 2) was used to make associative value *V* and latency data *L* comparable:

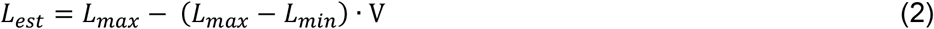

Where maximum latency is set to 5 and minimum latency is computed for each individual and is the minimum of a 100-trial moving average window. Therefore, the estimated latency is equal to 5 when associative value is 0 in the beginning and is equal to individual’s minimum latency when associative learning is 1, meaning subject has reached its best performance and learning is complete.

Finally, learning rate α is estimated for each animal by minimizing the sum of squared errors between latency data and the estimated latency.

### Electrophysiology

#### Recording procedures

Electrophysiology recordings of single unit spiking activity were conducted during both conditioning paradigms. Laboratory-made 8-channel electrode arrays (50μm diameter tungsten wire insulated with polyimide, California Fine Wire Company, Grover Beach, CA) were implanted in the VTA (AP −5.3, ML 0.8, DV −7.7) and SN (AP −5.2, ML 2.2, DV −7.4) under isoflurane anesthesia. All animals had one week to recover from surgery before the start of habituation. Recording began on the first day of conditioning training. During recordings, animals were connected via a field-effect transistor head-stage (Omnetics Connector Corp, Minneapolis, MN) to a lightweight cable and a rotating motorized commutator to allow unrestricted movement during recording. Spikes were amplified at 1000x gain, digitized at 40 kHz, and single-unit data was band-pass filtered at 300Hz. Single units were isolated in Kilosort (Allen et al., 2018) or Offline Sorter (Plexon) using a combination of manual and semi-automatic sorting techniques until each unit is well isolated in state space; minimum acceptable signal to noise ratio approximately 2.5:1. Neurons were not screened for specific physiological characteristics or response properties prior to recording.

#### Dopamine classification

Neurons were classified as putative dopamine based on wave form width greater than 1.2 ms and mean baseline firing rate slower than 12 Hz (Grace and Bunney, 1984; Schultz and Romo, 1987; Kim et al., 2016) consistent with the profile of optogenetically tagged dopamine neurons in our laboratory (Lohani et al., 2019). Additionally, a dopamine agonist drug study was conducted on a subset of subjects following the final recording session. After a 30 min baseline recording, animals were injected with 0.75 mg/kg apomorphine intraperitoneally and recorded for an additional 30 minutes (Grace and Bunney, 1984). Responsive units were defined through comparison of inter-spike interval distributions in a nonparametric Kolmogorov-Smirnov test (p<0.05). The direction of modulation after apomorphine (inhibited or excited) was determined by whether the pre- or post-injection distribution had a larger cumulative distribution function. Only neurons characterized as putative dopamine are used for firing rate analyses.

#### Spike Correlations

Spike correlations were computed by calculating the trial-by-trial correlation in spike counts between each pair of simultaneously recorded neurons as described previously (Kim et al., 2012). A Pearson’s correlation of spike counts was calculated in the 500 ms window following behavioral epochs of interest, consistent with firing rate analyses. Spike count correlations are sensitive to outliers, so we excluded any trial in which either unit firing rate was >3 standard deviations away from its mean baseline firing rate (Kohn and Smith, 2005; Ruff and Cohen, 2014). In order to preserve sufficient sample sizes, unit pairs were not grouped based on neurotransmitter content.

### Immunohistochemistry

#### Tissue preparation

All rats were anesthetized with chloral hydrate and perfused transcardially with phosphate buffered saline (PBS, pH 7.4, Sigma Aldrich) followed by 4% paraformaldehyde (PFA, Sigma Aldrich). Brains were extracted and post-fixed in 4% PFA overnight before being transferred and stored in a 20% sucrose solution at 4°C for cryoprotection. 40μm serial coronal sections were sliced using a cryostat and stored as free-floating sections in PBS + 0.05% sodium azide (NaN3).

#### Immunohistochemistry

Free-floating brain sections were blocked and permeabilized in a solution of PBS with 0.05% NaN3, 3% bovine serum albumin (BSA), 0.5% Triton-X, and 0.05% Tween-80 for two hours at room temperature. Sections were then incubated with a chicken polyclonal primary antibody against tyrosine hydroxylase (dilution 1:1000, Abcam ab76442) for 48 hours at 4°C. Sections were washed in a solution of PBS + 0.05% NaN3, 3% BSA, 0.1% Triton-X, and 0.01% Tween 80 before being incubated with an Alexa Fluor-conjugated goat anti-chicken secondary antibody (dilution 1:1000, Alexa Fluor 594, Abcam ab150176) for 48 hrs at 4°C. Sections were again washed before being mounted and cover-slipped with Vectashield hard-set mounting medium for fluorescence with DAPI (Vector Laboratories, Inc.).

#### Microscopy

A Zeiss Axiovert 200 microscope (Zeiss) with an Axiocam camera and Apotome II instrument with grid-based optical sectioning was used to visualize dopamine cells (tyrosine-hydroxylase labelled cells) on the red channel. Each image was acquired with the 20x objective and the Zen 2 software (Zeiss) generated a Z-stack scan series, consisting of 25 1-μm scans, resulting in three-dimensional images with a total volume of 120 × 120 × 25 μm^3^ per image stack. In all cases, four images from each brain were analyzed and averaged for each rat.

### Microdialysis

For microdialysis experiments, adolescent (PND 35–38) and adult (PND 65–70) rats were implanted with guide cannulas in both the medial area of the dorsal striatum (DS; AP= +1.6 mm ML= 2.2 mm from Bregma and DV= −4.0 mm from skull for adults and AP= +0.7 mm ML= 2.0 mm from Bregma and DV= −3.0 mm from skull for adolescents) and the nucleus accumbens (NA; AP= +1.2 mm ML= 1.1 mm from Bregma and DV= −6.0 mm from skull for adults and AP= +1.0 mm ML= 0.9 mm from Bregma and DV= −5.0 mm from skull for adolescents), plus a bipolar stimulating electrode in the VTA (AP= −5.3 mm ML= 0.9 mm from Bregma and DV = −8.3 mm from skull for adults and AP= −4.2 ML= −0.6 from Bregma and DV= −7.4 from skull for adolescents). One week after the surgery, microdialysis experiments on freely moving animals were conducted. Dialysis probes (CMA Microdialysis) with an active membrane length of 2 mm were inserted into the guide cannula and Ringers solution (in mM: 37.0 NaCl, 0.7 KCl, 0.25MgCl2, and 3.0 CaCl2) was perfused at flow rate of 2.0 μL/min. After 60 min of stabilization, dialysis samples (20 min each) were collected and immediately injected into an HPLC system with electrochemical detection of dopamine as described before (Adams and Moghaddam, 1998; Pehrson and Moghaddam, 2010). Once three consecutive stable baseline samples were observed, the electrical stimulation was delivered. The VTA was electrically stimulated for 20 minutes using one of two burst protocols: (a) the phasic burst stimulation (20 pulses at 100 Hz; pulse width = 1 ms, burst width = 200 ms, interburst interval (IBI) = 500 ms) or (b) the phasic sustained stimulation (100 pulses at 20 Hz; pulse width = 5 ms, burst width = 5 s, IBI = 10 s; Lohani et al., 2018; 2019). Microdialysis data were expressed as the percentage of baseline dopamine release, where baseline is defined as the mean of three consecutive samples obtained before the electrical stimulation. Motor behavior was measured during the microdialysis experiments by placing a stainless-steel frame with an array of infrared beams (Hamilton-Kinder, LLC, Poway, CA) outside the rats’ home-cage environment. Beam breaks were monitored over the entire course of the experiment using the Kinder Scientific Motor Monitor program. Locomotor activity data is expressed in terms of basic movements (total X/Y breaks) and values were pooled into 20-min bins corresponding to the collection of dialysis samples.

### Quantification and statistical analyses

#### Behavior and Electrophysiology

All analyses were performed in MATLAB (MathWorks, Nattick, MA) and R (https://www.r-project.org/). NeuroExplorer (NEX Technologies, Madison, AL) was used for preliminary analysis such as perievent rasters. For electrophysiological recordings, we conservatively classified neurons recorded in consecutive recording sessions as different units, despite any indications that the same unit were recorded serially. Unit firing rates for both behaviors were analyzed in 25 ms bins, and aligned with behavioral events. Baseline rate for individual units was determined using the average firing rate during the middle three s of the inter-trial interval. Isolated single unit data were analyzed with custom written Matlab functions. Statistical tests were performed using activity in 500 ms epoch windows, before, during and after the event of interest. Firing rate data collapsed across session was Z-score normalized, relative to baseline. Area under the curve was computed to assess group differences in firing rate. Group differences in physiology and behavior were assessed using mixed effect Analysis of variance (ANOVA) models. Dunnett’s Post hoc and Bonferroni corrected comparisons were used when appropriate.

#### Immunohistochemistry

IMARIS software (v.9.2.0; Bitplane) was used for image processing and quantification of the parameters of interest. Dopamine cells were visualized by tyrosine hydroxylase staining visible on the red channel, and IMARIS analysis modules were used for automated quantification of volume and number of dopamine cells within the VTA and SN. Statistical analyses were carried out in R (https://www.r-project.org/). Two-way ANOVAs were used to compare dopamine cell size and number across adult and adolescent rats in the VTA and SN.

#### Microdialysis

Microdialysis data were expressed as a percentage of baseline dopamine release, with baseline defined as the mean of three consecutive samples obtained before the electrical stimulation. For locomotor activity, data is expressed as the number of infrared beam breaks within a 20-min bin (which correspond with the 20-min microdialysis sample-collection). The statistical analysis of these dependent measures was conducted using two-factor repeated measures ANOVA with age as the between-subjects factor and time as the within-subjects factor.

#### Histology

After experiments were complete, rats were anesthetized and transcardially perfused with 0.9% saline, followed by 10% buffered formalin. Brains were stored in this formalin and transferred to 30% sucrose for at least 24 hours before brains were coronally sliced. Electrode and probe placement in the VTA, SN, DS and NA was confirmed for all animals who provided electrophysiological and microdialysis data.

## Notes

### Competing Interest Statement

The authors have declared no competing interest.

## References

Adams B, Moghaddam B (1998) Corticolimbic dopamine neurotransmission is temporally dissociated from the cognitive and locomotor effects of phencyclidine. Journal of Neuroscience 18:5545–5554.

Allen M, Chowdhury T, Wegener M, Moghaddam BJB (2018) Efficient sorting of single-unit activity from midbrain cells using KiloSort is as accurate as manual sorting.303479.

Averbeck BB, Costa VD (2017) Motivational neural circuits underlying reinforcement learning. Nature Neuroscience 20:505–512.

Barker JM, Corbit LH, Robinson DL, Gremel CM, Gonzales RA, Chandler LJ (2015) Corticostriatal circuitry and habitual ethanol seeking. Alcohol 49:817–824.

Berridge KC (2004) Motivation concepts in behavioral neuroscience. Physiology & Behavior 81:179–209.

Cardinal RN, Parkinson JA, Hall J, Everitt BJ (2002) Emotion and motivation: the role of the amygdala, ventral striatum, and prefrontal cortex. Neuroscience & Biobehavioral Reviews 26:321–352.

Casey B (2015) Beyond simple models of self-control to circuit-based accounts of adolescent behavior. Annual review of psychology 66:295–319.

Coddington LT, Dudman JTJNn (2018) The timing of action determines reward prediction signals in identified midbrain dopamine neurons. 21:1563.

Cohen JY, Haesler S, Vong L, Lowell BB, Uchida N (2012) Neuron-type-specific signals for reward and punishment in the ventral tegmental area. nature 482:85.

Cohen MR, Kohn A (2011) Measuring and interpreting neuronal correlations. Nature neuroscience 14:811.

Corbit LH, Janak PH (2010) Posterior dorsomedial striatum is critical for both selective instrumental and Pavlovian reward learning. European Journal of Neuroscience 31:1312–1321.

Corbit LH, Balleine BW (2016) Learning and Motivational Processes Contributing to Pavlovian– Instrumental Transfer and Their Neural Bases: Dopamine and Beyond. In: Behavioral Neuroscience of Motivation (Simpson EH, Balsam PD, eds), pp 259–289. Cham: Springer International Publishing.

Dalley JW, Chudasama Y, Theobald DE, Pettifer CL, Fletcher CM, Robbins TW (2002) Nucleus accumbens dopamine and discriminated approach learning: interactive effects of 6-hydroxydopamine lesions and systemic apomorphine administration. Psychopharmacology 161:425–433.

Danks D (2003) Equilibria of the Rescorla–Wagner model. Journal of Mathematical Psychology 47:109–121.

Day JJ, Carelli RM (2007) The nucleus accumbens and Pavlovian reward learning. The Neuroscientist 13:148–159.

Dayan P, Balleine BW (2002) Reward, motivation, and reinforcement learning. Neuron 36:285–298.

Dickinson A (1981) Conditioning and associative learning. British Medical Bulletin 37:165–168.

Ernst M, Luciana M (2015) Neuroimaging of the dopamine/reward system in adolescent drug use. CNS spectrums 20:427.

Ernst M, Daniele T, Frantz K (2011) New perspectives on adolescent motivated behavior: attention and conditioning. Developmental cognitive neuroscience 1:377–389.

Fanselow MS, Wassum KM (2015) The Origins and Organization of Vertebrate Pavlovian Conditioning. Cold Spring Harb Perspect Biol 8:a021717.

Flagel SB, Clark JJ, Robinson TE, Mayo L, Czuj A, Willuhn I, Akers CA, Clinton SM, Phillips PE, Akil H (2011) A selective role for dopamine in stimulus-reward learning. Nature 469:53–57.

Gopnik A, O’Grady S, Lucas C, Griffiths T, Wente A, Bridgers S, Aboody R, Fung H, Dahl R Submission PDF. PNAS 67:68.

Grace AA, Bunney BS (1984) The control of firing pattern in nigral dopamine neurons: single spike firing. Journal of neuroscience 4:2866–2876.

Gremel CM, Costa RM (2013) Orbitofrontal and striatal circuits dynamically encode the shift between goal-directed and habitual actions. Nature communications 4:2264.

Guyenet P, Aghajanian G (1978) Antidromic identification of dopaminergic and other output neurons of the rat substantia nigra. Brain research 150:69–84.

Haber SN (2016) Corticostriatal circuitry. Dialogues in clinical neuroscience 18:7.

Haruno M, Kawato M (2006) Different neural correlates of reward expectation and reward expectation error in the putamen and caudate nucleus during stimulus-action-reward association learning. Journal of neurophysiology 95:948–959.

Hauser TU, Will GJ, Dubois M, Dolan RJ (2019) Annual research review: developmental computational psychiatry. Journal of Child psychology and Psychiatry 60:412–426.

Hughes RN, Bakhurin KI, Petter EA, Watson GD, Kim N, Friedman AD, Yin HH (2020) Ventral tegmental dopamine neurons control the impulse vector during motivated behavior. Current Biology.

Huttenlocher PR (1979) Synaptic density in human frontal cortex — Developmental changes and effects of aging. Brain Research 163:195–205.

Keiflin R, Pribut HJ, Shah NB, Janak PH (2019) Ventral tegmental dopamine neurons participate in reward identity predictions. 29:93-103. e103.

Kim Y, Wood J, Moghaddam B (2012) Coordinated activity of ventral tegmental neurons adapts to appetitive and aversive learning. PloS one 7.

Kim Y, Simon NW, Wood J, Moghaddam B (2016) Reward Anticipation Is Encoded Differently by Adolescent Ventral Tegmental Area Neurons. Biological Psychiatry 79:878–886.

Kohn A, Smith MA (2005) Stimulus dependence of neuronal correlation in primary visual cortex of the macaque. Journal of Neuroscience 25:3661–3673.

Larsen B, Luna B (2018) Adolescence as a neurobiological critical period for the development of higher-order cognition. Neuroscience & Biobehavioral Reviews 94:179–195.

Lebel C, Walker L, Leemans A, Phillips L, Beaulieu C (2008) Microstructural maturation of the human brain from childhood to adulthood. Neuroimage 40:1044–1055.

Lex B, Hauber W (2010) The role of dopamine in the prelimbic cortex and the dorsomedial striatum in instrumental conditioning. Cerebral cortex 20:873–883.

Lohani S, Martig AK, Deisseroth K, Witten IB, Moghaddam B (2019) Dopamine modulation of prefrontal cortex activity is manifold and operates at multiple temporal and spatial scales. Cell reports 27:99-114. e116.

Luciana M, Collins PF (2012) Incentive motivation, cognitive control, and the adolescent brain: Is it time for a paradigm shift? Child development perspectives 6:392–399.

Luna B, Marek S, Larsen B, Tervo-Clemmens B, Chahal R (2015) An integrative model of the maturation of cognitive control. Annual review of neuroscience 38:151–170.

Maren S (2001) Neurobiology of Pavlovian fear conditioning. Annu Rev Neurosci 24:897–931.

Marshall AT, Munson CN, Maidment NT, Ostlund SB (2020) Reward-predictive cues elicit excessive reward seeking in adolescent rats. Developmental cognitive neuroscience 45:100838.

McDannald MA, Saddoris MP, Gallagher M, Holland PC (2005) Lesions of orbitofrontal cortex impair rats’ differential outcome expectancy learning but not conditioned stimulus-potentiated feeding. Journal of Neuroscience 25:4626–4632.

Mohebi A, Pettibone JR, Hamid AA, Wong J-MT, Vinson LT, Patriarchi T, Tian L, Kennedy RT, Berke JD (2019) Dissociable dopamine dynamics for learning and motivation. Nature 570:65–70.

Naneix F, Marchand AR, Di Scala G, Pape J-R, Coutureau E (2012) Parallel maturation of goal-directed behavior and dopaminergic systems during adolescence. Journal of Neuroscience 32:16223–16232.

O’Doherty J, Dayan P, Schultz J, Deichmann R, Friston K, Dolan RJ (2004) Dissociable roles of ventral and dorsal striatum in instrumental conditioning. science 304:452–454.

O’Doherty JP (2016) Multiple Systems for the Motivational Control of Behavior and Associated Neural Substrates in Humans. Curr Top Behav Neurosci 27:291–312.

Parkinson JA, Dalley JW, Cardinal RN, Bamford A, Fehnert B, Lachenal G, Rudarakanchana N, Halkerston KM, Robbins TW, Everitt BJ (2002) Nucleus accumbens dopamine depletion impairs both acquisition and performance of appetitive Pavlovian approach behaviour: implications for mesoaccumbens dopamine function. Behavioural Brain Research 137:149–163.

Peak J, Hart G, Balleine BW (2019) From learning to action: the integration of dorsal striatal input and output pathways in instrumental conditioning. European Journal of Neuroscience 49:658–671.

Pehrson AL, Moghaddam B (2010) Impact of metabotropic glutamate 2/3 receptor stimulation on activated dopamine release and locomotion. Psychopharmacology 211:443–455.

Pessiglione M, Seymour B, Flandin G, Dolan RJ, Frith CD (2006) Dopamine-dependent prediction errors underpin reward-seeking behaviour in humans. Nature 442:1042–1045.

Pitts EG, Stowe TA, Christensen BA, Ferris MJ (2020) Comparing dopamine release, uptake, and D2 autoreceptor function across the ventromedial to dorsolateral striatum in adolescent and adult male and female rats. Neuropharmacology 175:108163.

Robbins TW, Everitt BJ (1996) Neurobehavioural mechanisms of reward and motivation. Current opinion in neurobiology 6:228–236.

Robinson DL, Zitzman DL, Smith KJ, Spear LP (2011) Fast dopamine release events in the nucleus accumbens of early adolescent rats. Neuroscience 176:296–307.

Ruff DA, Cohen MR (2014) Attention can either increase or decrease spike count correlations in visual cortex. Nature neuroscience 17:1591–1597.

Salamone JD, Correa M, Yang J-H, Rotolo R, Presby R (2018) Dopamine, effort-based choice, and behavioral economics: basic and translational research. Frontiers in behavioral neuroscience 12:52.

Saunders BT, Richard JM, Margolis EB, Janak PH (2018) Dopamine neurons create Pavlovian conditioned stimuli with circuit-defined motivational properties. Nature neuroscience 21:1072–1083.

Schultz W (1998) Predictive Reward Signal of Dopamine Neurons. Journal of Neurophysiology 80:1–27.

Schultz W (2010) Multiple functions of dopamine neurons. F1000 biology reports 2.

Schultz W, Romo R (1987) Responses of nigrostriatal dopamine neurons to high-intensity somatosensory stimulation in the anesthetized monkey. Journal of neurophysiology 57:201–217.

Serlin H, Torregrossa MM (2015) Adolescent rats are resistant to forming ethanol seeking habits. Developmental cognitive neuroscience 16:183–190.

Shiflett MW, Balleine BW (2011) Molecular substrates of action control in cortico-striatal circuits. Prog Neurobiol 95:1–13.

Simon NW, Moghaddam B (2015) Neural processing of reward in adolescent rodents. Developmental cognitive neuroscience 11:145–154.

Spear LP (2000) The adolescent brain and age-related behavioral manifestations. Neuroscience & Biobehavioral Reviews 24:417–463.

Stauffer WR, Lak A, Yang A, Borel M, Paulsen O, Boyden ES, Schultz W (2016) Dopamine neuron-specific optogenetic stimulation in rhesus macaques. Cell 166:1564-1571. e1566.

Sturman DA, Moghaddam B (2011) Reduced neuronal inhibition and coordination of adolescent prefrontal cortex during motivated behavior. The Journal of Neuroscience 31:1471–1478.

Sturman DA, Moghaddam B (2012a) Striatum processes reward differently in adolescents versus adults. Proc Natl Acad Sci U S A 109:1719–1724.

Sturman DA, Moghaddam B (2012b) Striatum processes reward differently in adolescents versus adults. Proceedings of the National Academy of Sciences 109:1719–1724.

Towner TT, Fager M, Spear LP (2020) Adolescent but not adult Sprague-Dawley rats display goaldirected responding after reward devaluation. Developmental psychobiology 62:368–379.

van Zessen R, Flores-Dourojeanni JP, Eekel T, van den Reijen S, Lodder B, Omrani A, Smidt MP, Ramakers GM, van der Plasse G, Stuber GD (2021) Cue and reward evoked dopamine activity is necessary for maintaining learned Pavlovian associations. Journal of Neuroscience 41:5004–5014.

Yin HH, Ostlund SB, Knowlton BJ, Balleine BWJEJoN (2005) The role of the dorsomedial striatum in instrumental conditioning. 22:513–523.

